# Emergence of Multidrug-Resistant Uropathogens harboring ESBL, Carbapenem, Aminoglycosides and AmpC resistant genes from Northern India

**DOI:** 10.1101/375501

**Authors:** Varsha Rani Gajamer, Amitabha Bhattacharjee, Deepjyoti Paul, Birson Ingti, Arunabha Sarkar, Jyotsna Kapil, Ashish Kumar Singh, Nilu Pradhan, Hare Krishna Tiwari

## Abstract

Extended-spectrum β-lactamase (ESBL) producing bacteria acts as a serious threat, and its co-existence with other antibiotic resistant gene makes the clinical scenario worse nowadays. Therefore in this study, we investigated the occurrence of ESBL genes coexisting with carbapenem, AmpC and aminoglycoside resistance gene in uropathogens. Out of 1516 urine samples, 454 showed significant bacteriuria with a prevalence rate of 29.94 %. *Escherichia coli* (n=340) were found to be the most predominant uropathogen followed by *Klebsiella pneumoniae* (n=92), *Pseudomonas aeruginosa* (n=10) and *Proteus mirabilis* (n=9). Among the total uropathogens, sixty-three ESBL-producers were identified which included *bla*_CTX-M-15_ (n=32), followed by *bla*_CTX-M-15_ + *bla*_OXA-2_ (n=15), *bla*_CTX-M-15_ + *bla*_OXA-2_ + *bla*_TEM_ (n=6), *bla*_OXA-2_ (n=5), *bla* _OXA-2_ + *bla* _SHV-76_ (n=1), *bla* _TEM_+_SHV-76_ (n= 1) and *bla* _TEM_ (n=1). All ESBL genes were found on plasmid incompatibility types: HI1, I1, FIA+FIB, FIA and Y and were horizontally transferable. Among 63 ESBL-producers, 59 isolates harboured carbapenem-resistant genes which included *bla*_NDM-5_ (n=48), *bla*_NDM-5_ + *bla*_OXA-48_ (n=5), *bla*_NDM-5_ + *bla*_IMP_ (n=5) and *bla*_NDM-5_ + *bla*_IMP_ + *bla*_VIM_ (n=1). The ESBL producing uropathogens also harbored 16S rRNA methylase genes which included *rmtB* (n=9), *rmtA* (n=4), *rmtC* (n=1) and *ArmA* (n=1) followed by AmpC genes which includes CIT (n=8) and DHA-1 (n=1) genes. Imipenem and gentamicin were found to be more effective. We speculating, this is the first report showing the prevalence of multidrug-resistant uropathogens in this area demanding regular surveillance for such resistance mechanisms which will be useful for health personnel to treat ESBL infection and its co-existence with another antibiotic resistance gene.

## INTRODUCTION

The rise in the prevalence of antibiotic resistance and the lack of new antibiotic drug development has gradually reduced the treatment options for bacterial infections (1). Massive use of antibiotics plays a crucial role in the emergence of antibiotic resistance among gram-negative bacteria worldwide (2). Due to the huge selective pressure of antibiotics, and the presence of various ESBLs in different countries ß-lactamases are remarkably diversified. Since ESBL genes are mostly plasmid mediated, it may also carry genes encoding resistance to other class of antibiotics, such as aminoglycosides, macrolides, chloramphenicol, quinolone or carbapenems. Due to the presence of the multiple resistance genes encoded in the plasmids, treatment options become restricted for ESBL producing bacteria (3).

The emergence and rapid distribution of ESBL producing bacteria which are capable of hydrolyzing penicillins, broad-spectrum cephalosporins, and monobactams, also harbor resistance genes for other antibiotics, thus making carbapenem limiting treatment options for infections over the last couple of decades (4). Carbapenemase-producing *E. coli* are of major clinical concern (5), It has been reported that carbapenem-resistant Enterobacteriaceae causing mortality by up to 50% of patients who acquire bloodstream infections (6). As a consequence of this, increased utilization of carbapenems has led to the emergence of isolates with resistance genes that code for carbapenemases.

AmpC harboring strains are challenging as it confers resistance to broad-spectrum cephalosporins which may further limit treatment option when expressed to higher levels (7). Moreover, aminoglycosides are commonly used in combination with the ß-lactam group of antibiotics for the treatment of severe infections in hospital patients. However, the bacterial population has developed various mechanisms of resistance and in a little while, the therapeutic use of this drug will be inadequate (8). Infections due to ESBL producers with the capacity of antibiotic resistance with other precious antibiotics like carbapenem, and aminoglycosides makes the treatment challenging. Therefore, the timely detection of such pathogens is always required to manage the infections with the judicial use of antibiotics.

Therefore, the present work was conducted, to study the emergence of ESBL genes in uropathogens, their co-existence with carbapenem-resistant genes followed by AmpC and Aminoglycoside resistant genes in Northern India.

## MATERIAL AND METHODS

### Sample collection and Isolation

The present study was conducted from May 2014 to September 2016 in the Department of Microbiology, Sikkim University, Sikkim, India. A total of 1516 non-duplicate urine samples were collected from UTI suspected female patients of age group18 to 48. The samples were collected from inpatient and out-patient departments of tertiary hospitals namely Sikkim Manipal Institute of Medical Sciences,Gangtok, Sikkim and Neotia Get Wel hospital Siliguri, West BengalThe standard microbiological techniques were used for the collection, transportation, and processing of clean-catch mid-stream samples. The uropathogens were isolated on Cystine Lactose Electrolyte Deficient Agar (CLED), Hi chrome UTI agar and Mac Conkey agar plate by the semi-quantitative method. Specimens yielding more than or equal to 10^5^cfu/ml of urine were interpreted as significant bacteriuria (9).

### Identification of uropathogens

All the isolates were identified on the basis of gram staining, colony morphology and standard biochemical tests. The representative strains were further identified by Vitek 2 instruments (VITEK 2 compact, Biomerieux, Germany) and further confirmed by 16s rDNA sequencing.

### Antibiotic Susceptibility Testing

Antibiotic susceptibility testing was performed by Kirby-Bauer disk diffusion method on the Muller Hinton Agar (MHA) as per Clinical Laboratory Standards Institute (CLSI,2011) guidelines to determine the drug resistance pattern of different isolated uropathogens (10).

### Phenotypic detection of ESBLs

The screening test of the isolates was done using five antibiotics namely cefotaxime, ceftazidime, ceftriaxone, aztreonam at 1μg/ml and cefpodoxime 4 μg/ml (1 μg/ml for *Proteus mirabilis*) in Mueller Hinton Agar by agar dilution method. The isolates that showed growth in any of these antibiotic containing medium was suspected to be ESBL producer and were subjected to a confirmatory test (10).

### Confirmatory tests for ESBL production –

Isolates considered to be positive for ESBL production by the screening test were subjected to the Phenotypic confirmatory test using ESBL kit (Himedia, Mumbai) consisting of ceftazidime (30 μg) (CAZ), ceftazidime + clavulanic acid (30/10 μg) (CAC), cefotaxime (30 μg) (CTX) and cefotaxime + clavulanic acid (30/10μg) (CEC) (10).

### Genotypic characterization of resistant genes

#### DNA Extraction

Total DNA from bacterial samples was collected by the boiling method. The organism was cultured in 5ml Luria Bertani broth. One ml of culture was added to an Eppendorf tube and centrifuged at 10,000 rpm for 5 minutes followed by supernatant was discarded and pellets were resuspended in sterile distilled water. The culture was heated/boiled at 85°C for 20 mins. The lysis cell was centrifuged at (10,000 rpm for 10 mins). The supernatant containing DNA was collected (11).

##### ESBL genes

ESBL genes were detected by multiplex PCR (BioRad, USA) in a total volume of 25 μl containing 23.5 master mix and 1.5 μl of template DNA. For amplification and characterization of *bla*_ESBL_, a set of eight primers were used namely: *bla*_TEM_, *bla*_CTX-M_, *bla*_SHV_, *bla*_OXA-2_, *bla*_OXA-1O_, *bla*_PER_, *bla*_GES_ and *bla*_VEB_ (Table 1). Reactions were run under the following conditions: initial denaturation at 94°C for five min, 33 cycles of 94 °C for 35 sec, 51°C for one min, 72°C for one min and the final extension at 72°C for seven min^8^ PCR products were separated by gel electrophoresis on 1 % agarose gel.

**Table 1:**
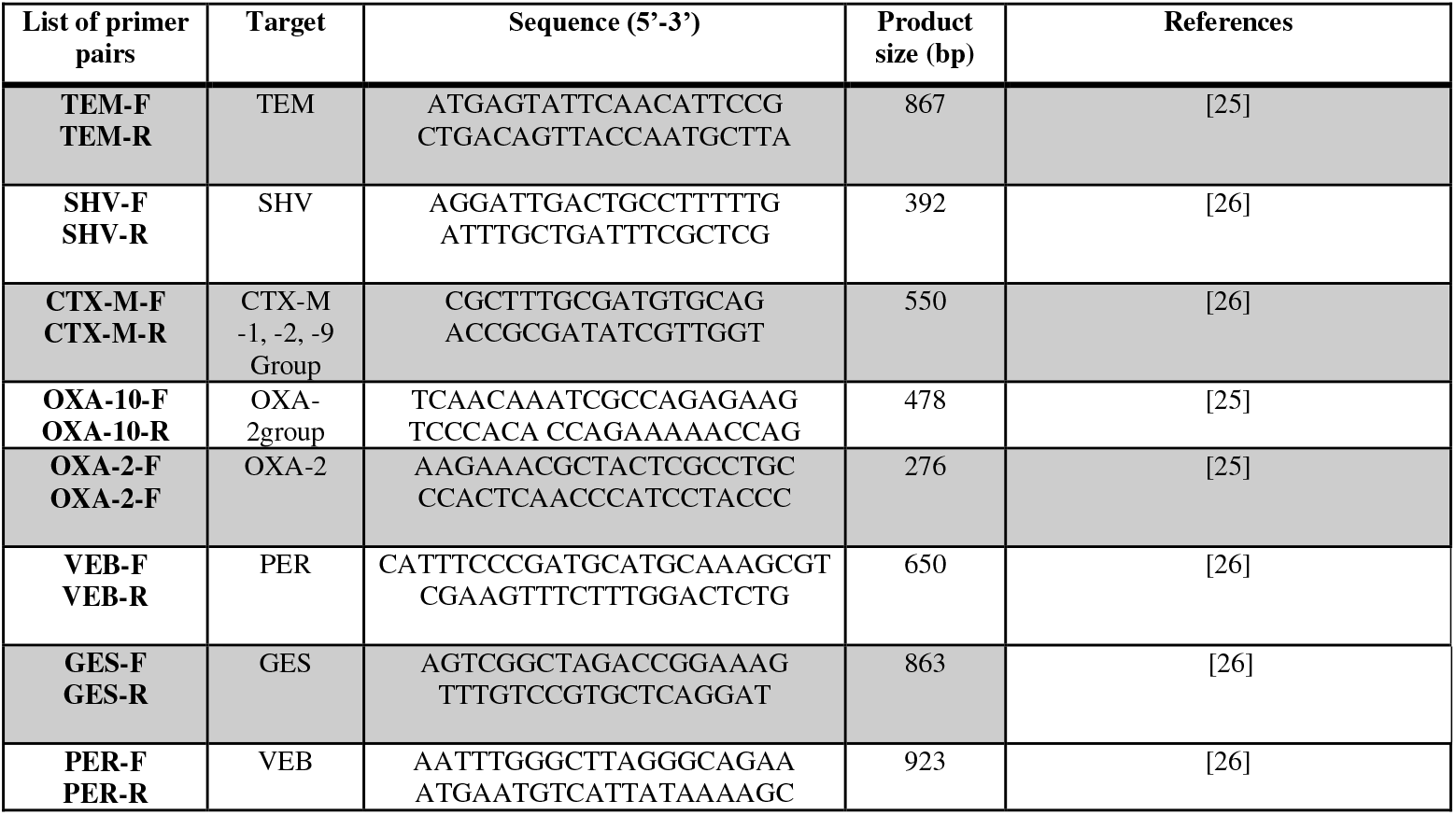
Detailed information of primers used in multiplex PCR for detection of *bla* genes in ESBL producers

##### Aminoglycoside resistant genes

The emergence of isolates resistant to all clinically important aminoglycosides related to the production of 16S rRNA methylases is worrisome. Molecular characterization of aminoglycoside-resistant gene was performed by a multiplex PCR targeting six 16S rRNA methyltransferase genes namely *armA, rmtA, rmtB, rmtC, rmtD*, and *npmA* (12) (Table 2). These genes were characterized by two multiplex PCR assays and further confirmed by a simplex PCR. Reactions were performed under the following conditions; initial denaturation at 94°C for 5 min, 34 cycles of 94°C for 30 s, 52°C for 40 s, and 72°C for 1 min 20 s, and a final extension at 72°C for 7 min. The amplified products were further sequenced to confirm the presence of the resistant genes.

**Table 2:**
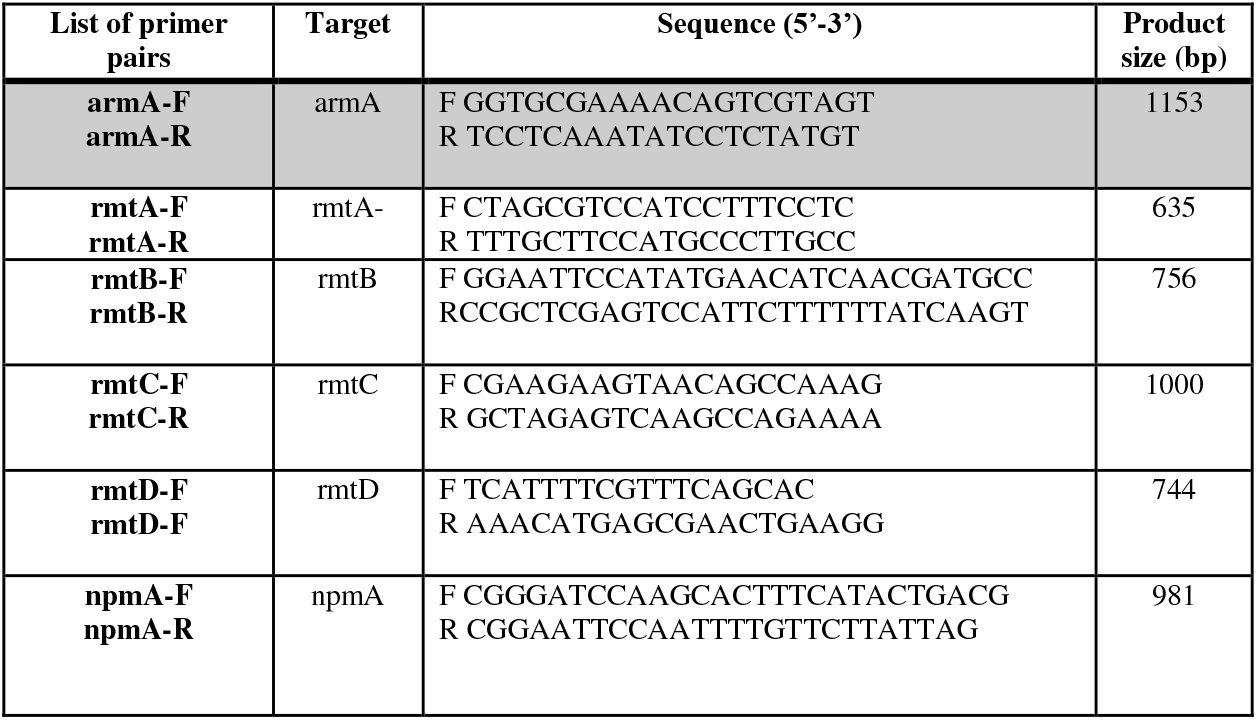
Detailed information of primers used in multiplex PCR for detection of 16S rRNA methyltransferase genes [10]

##### AmpC gene

Multiplex PCR was performed targeting all the AmpC genes by using a pair of primers as listed in Table 3. Isolates were investigated for the presence of other AmpC gene families namely; DHA, CIT, ACC, FOX and EBC (13). PCR amplification was performed using 30 μl of total reaction volume. Reactions were run under the following conditions: initial denaturation at 95 °C for 2 min, 34 cycles of 95 °C for 15 s, 51 °C for 1 min, 72 °C for 1 min and a final extension at 72 °C for 7 min.

**Table 3:**
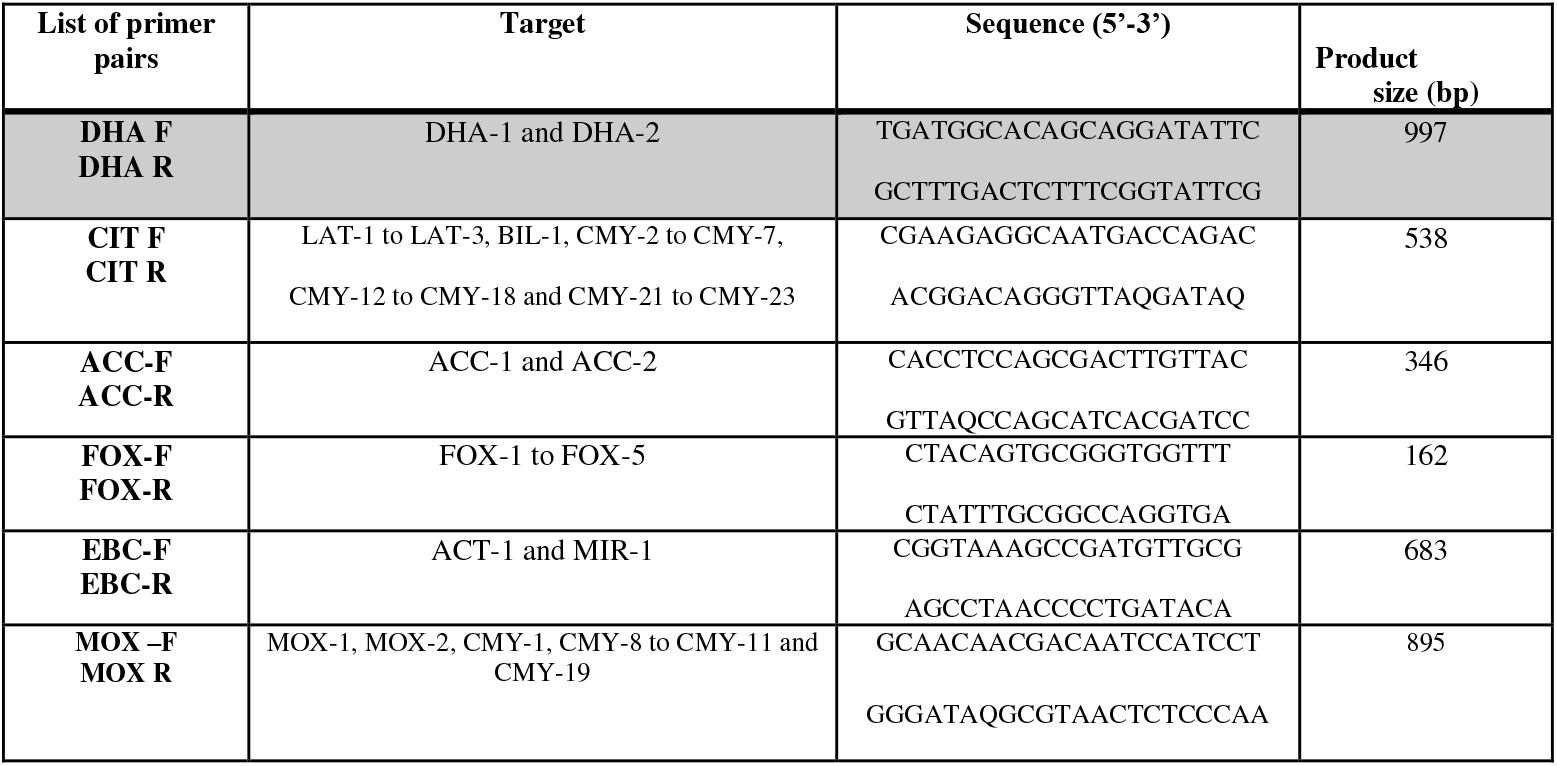
Detailed information of primers used in multiplex PCR for detection of AmpC resistant genes [11]

**Table 4:**
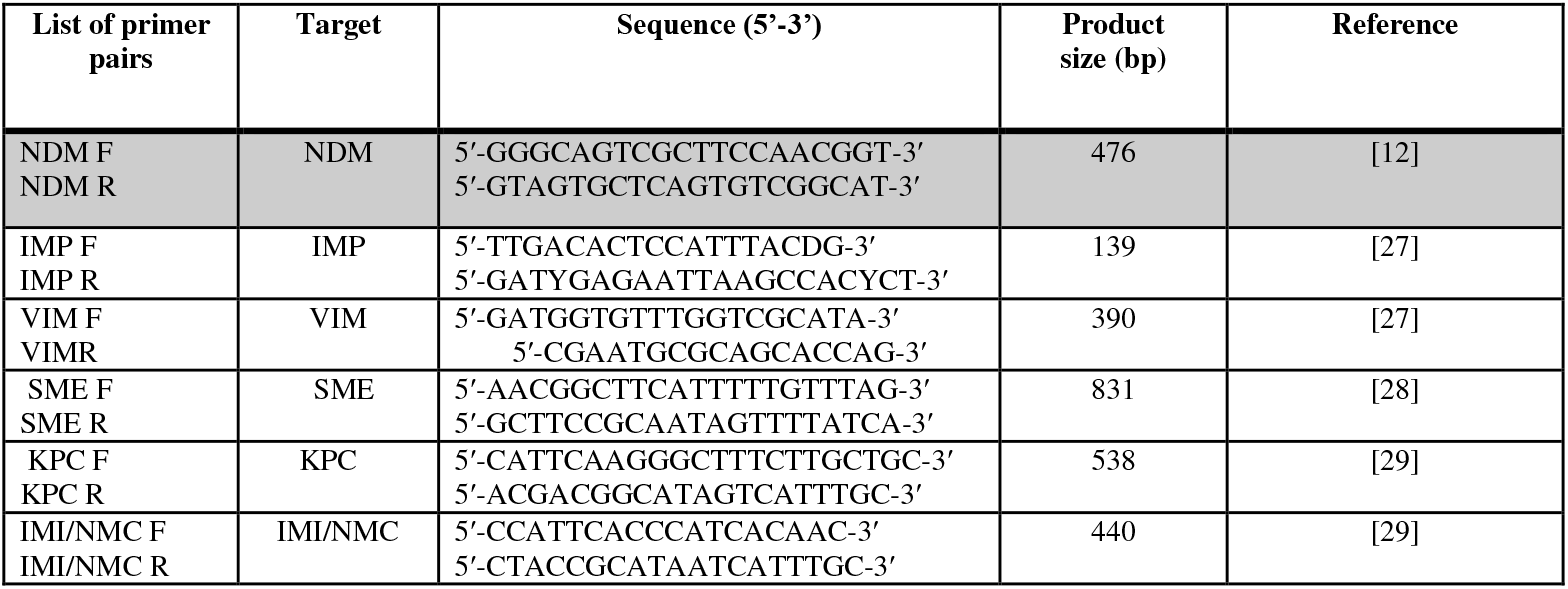
Detailed information of primers used in multiplex PCR for detection of Carbapenem resistant genes

##### Carbapenem-resistant gene

Carbapenem-resistant genes were detected by multiplex PCR (BioRad, USA) in a total volume of 30 μl. For amplification and characterization of the carbapenem-resistant gene, a set of nine primers were used namely *bla*_SME_, *bla*_IMI/NMC_, *bla*_KPC_, *bla*_NDM_, *bla*_VIM_, *bla*_IMP_, and *bla*_OXA-23_, *bla*_OXA-48_, *bla*_OXA-58_ (14). Reactions were performed under the following conditions: Initial denaturation at 94°C for 10 min; 30 cycles of 94°C for 40 sec, 55°C for 40 sec and 72°C for one min; and a final elongation step at 72°C for seven min. PCR products were separated by gel electrophoresis on 1 % agarose gel.

#### DNA sequence analysis

Total DNA from representative strains of each gene was prepared and purified by procedures of Upadhaya et al., 2015. (11). Sequencing was performed to identify specific ESBL, AmpC, Carbapenem, resistant genes. The DNA was sequenced using the dideoxynucleotide chain termination method at Sci genome, Kakkanad, Cochin, India. The ABI sequence files were assembled, and contigs were prepared using Codon Code aligner software (CodonCode Aligner 7.0.1.).

Nucleotide sequence similarity searches were performed using the National Centre for Biotechnology Information (NCBI) Basic Local Alignment Search Tool (BLAST) server on the GenBank database. (15).

#### Plasmid stability test of isolates

Plasmid stability of all *bla* producers as well as their transformants was analyzed by serial passages method for 110 consecutive days at 1:1000 dilutions in Luria-Bertani broth (Hi-Media, Mumbai, India) without antibiotic pressure (15). PCR assay was performed to check the presence of *bla* genes in the isolates after each passage.

#### Plasmid preparation, genetic transferability and incompatibility typing

Bacterial isolates harbouring ESBL genes were cultured in Luria-Bertani broth (Hi-Media, Mumbai, India) containing 0.25 μg/ml of cefoxitin. After an overnight incubation, plasmids were extracted by QIAprep Spin Miniprep Kit (Qiagen, Germany). Plasmids of *bla* genes were subjected to transformation by heat shock method. *E*.JM107 was used as a recipient. Transformants were selected on Luria-Bertani agar with 0.25 μg/ml of cefoxitin, which were then confirmed both by phenotypic as well as by genotypic method. The plasmids were classified by PCR based replicon typing(15), a total of 18 different replicon types namely FIA, FIB, FIC, HI1, HI2, I1/Iγ, L/M, N, P, W, T, A/C, K, B/O, X, Y, *F* and FIIA were targeted using 5 multiplex and 3 simplex PCR [15].

## RESULTS

Among the 1516 urine samples collected from female patients suspected to have UTI, 454 showed significant growth (significant bacteriuria) of a single type of microorganism with a prevalence rate of 29.94 %. Among 454 isolates, *E. coli* (n=340) were found to be the most predominant uropathogen followed by *Klebsiella pneumoniae* (n=92), *Pseudomonas aeruginosa* (n=10) and *Proteus mirabilis* (n=9).

Antibiotic susceptibility test revealed that 60-90% of the isolates showed resistance to ampicillin. Imipenem and gentamicin were found to be more effective (Fig:1-4). Among 86 isolates that were phenotypically confirmed as ESBL producers, 63 (n=*E.coli*-47, n= *K. pneumoniae*- 16) isolates showed the presence of β-lactamase genes by multiplex PCR. Sanger sequencing confirmed the occurrence of ESBL genes and also revealed its different variants. Four different ESBL gene variants were detected. *bla*_CTX-M-15_ was found to be the more prevalent and some genes were also found in combination like *bla*_CTX-M^−^15_ + *bla*_OXA-2_, *bla*_CTX-M-15_ + *bla*_OXA-2_ + *bla*_TEM_, *bla*_OXA-2_ + *bla*_SHV-76_ and *bla* _TEM_+_SHV^−^76_. Among the 63 ESBL–producing strains, 59 isolates harboured carbapenem-resistant genes which included *bla*_NDM-5_, *bla*_NDM-5_ + *bla*_OXA-48_, and *bla*_NDM-5_ + *bla*_IMP_ + *bla*_VIM_ and *bla*_NDM-5_ + *bla*_IMP_ (Fig:5-12).

**Fig 1:**
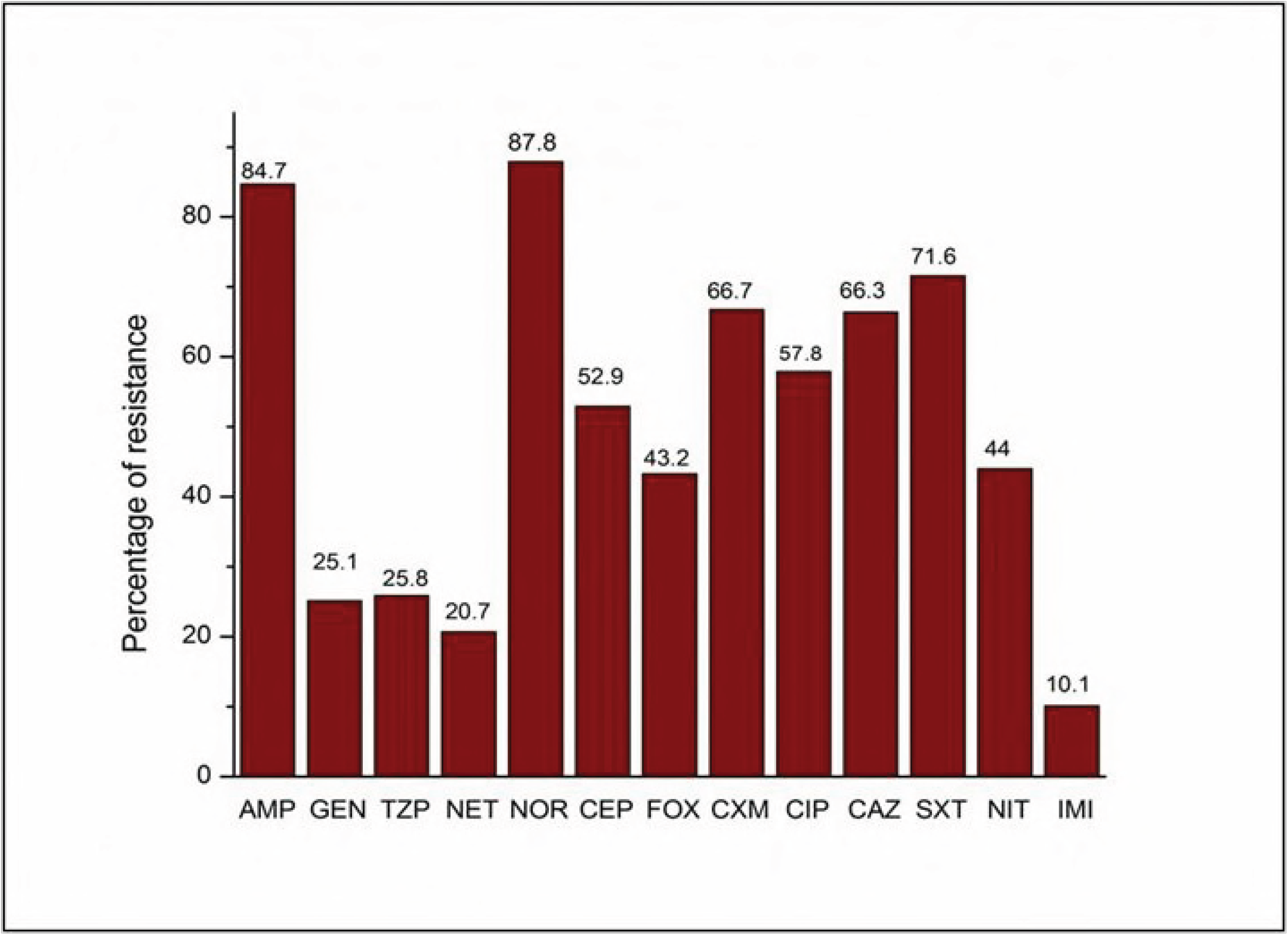
Resistance pattern of *Escherechia coli* against commonly used antibiotics

**Fig 2:**
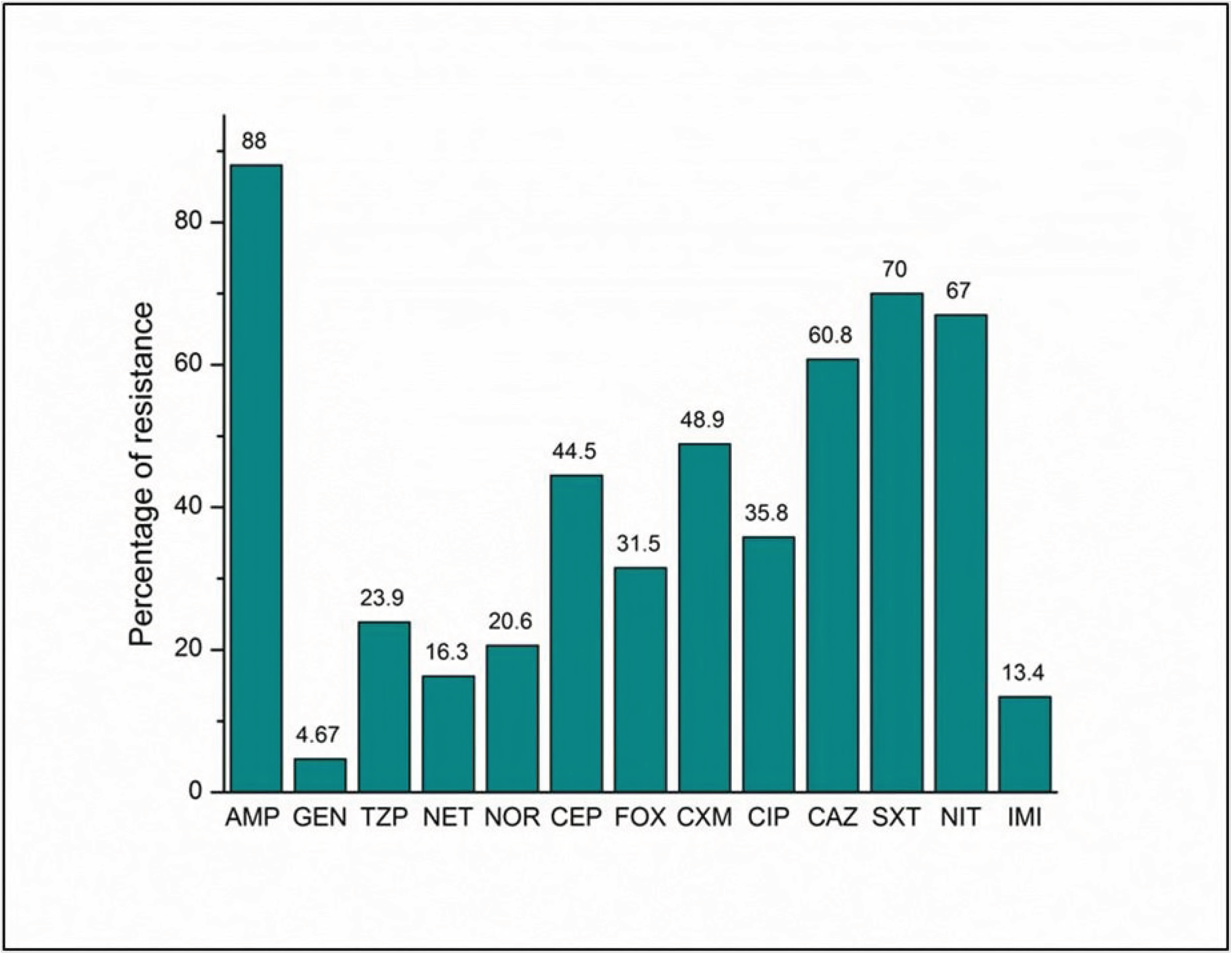
Resistance pattern of *Klebsiella pneumoniae* against commonly used antibiotics

**Fig 3:**
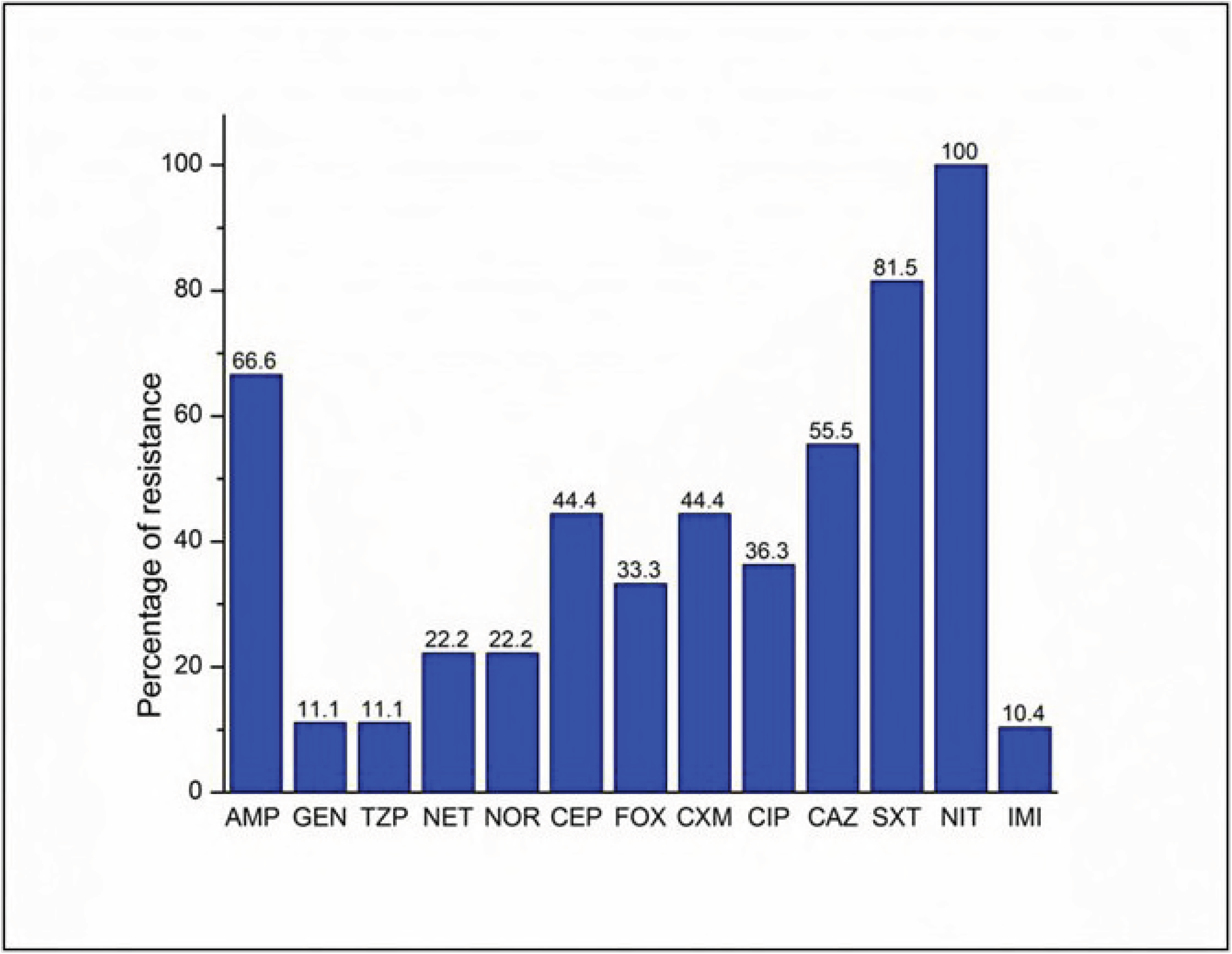
Resistance pattern of *Proteus mirabilis* against commonly used antibiotics

**Fig 4:**
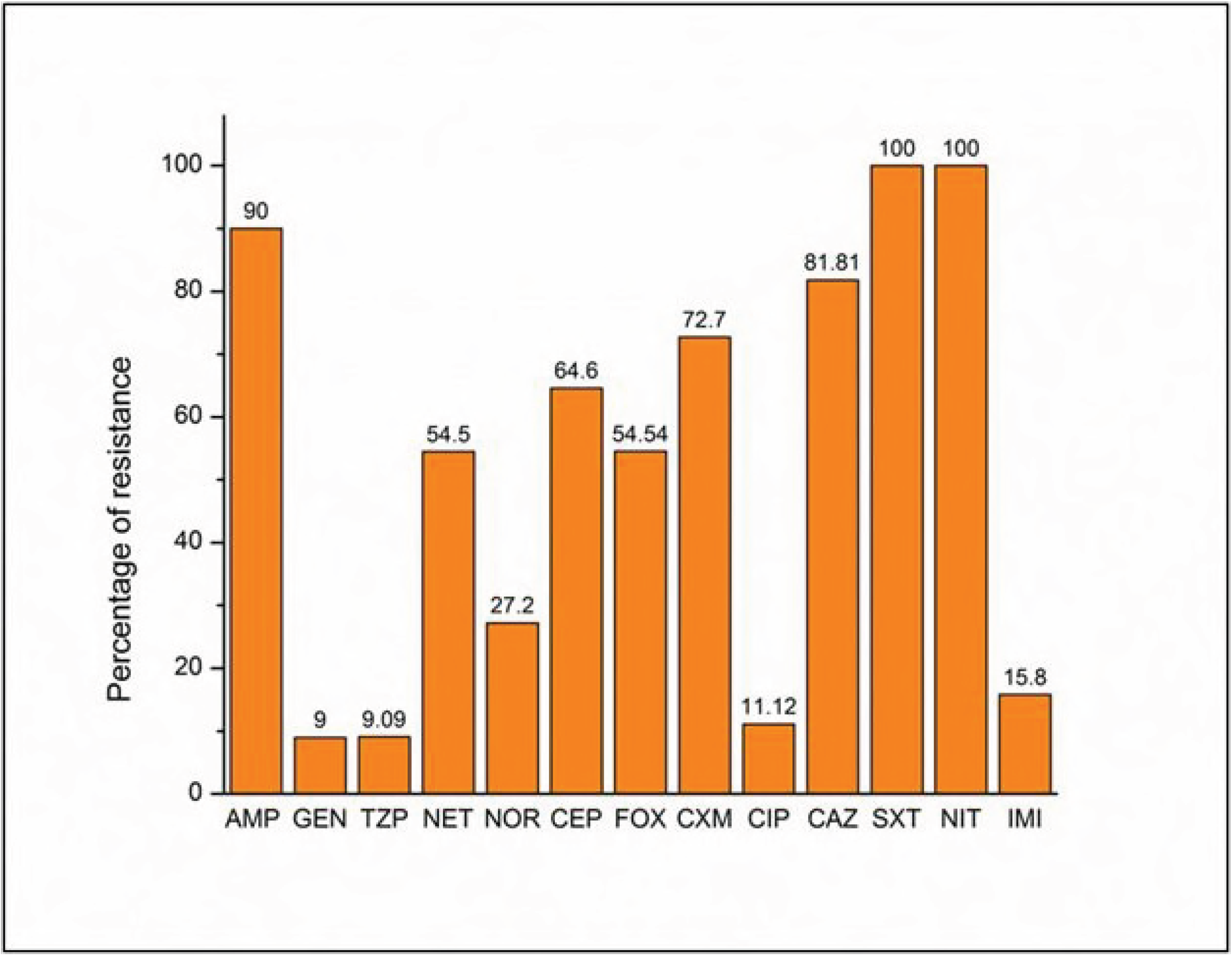
Resistance pattern of *Pseudomonas aeruginosa* against commonly used antibiotics

**Fig 5:**
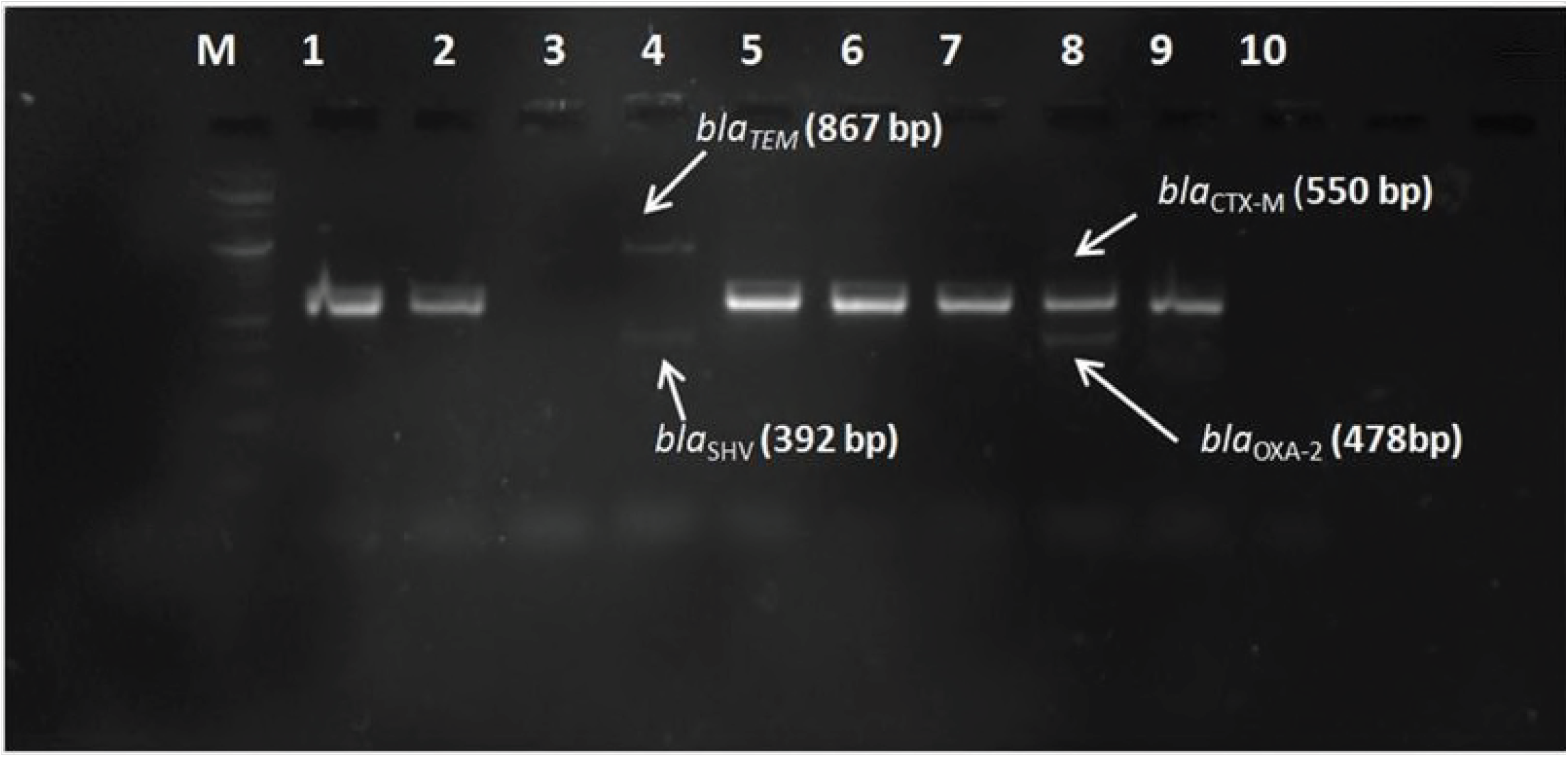
Agarose gel showing PCR amplified products of ESBL gene. Lane M-100bp DNA ladder. Lane 1-*bla*_CTX-M_, Lane 2 -*bla*_CTX-M_, Lane 3-No band, Lane 4- *bla*_TEM+SHV_, Lane 5-*bla*_CTX-M_, Lane 6- *bla*_CTX-_, Lane 7- *bla*_CTX-M_, Lane 8- *bla*_CTX-M+OXA-2_, Lane 9- +ve control (*bla*_CTX-M_), Lane 10- ve control.

**Fig 6:**
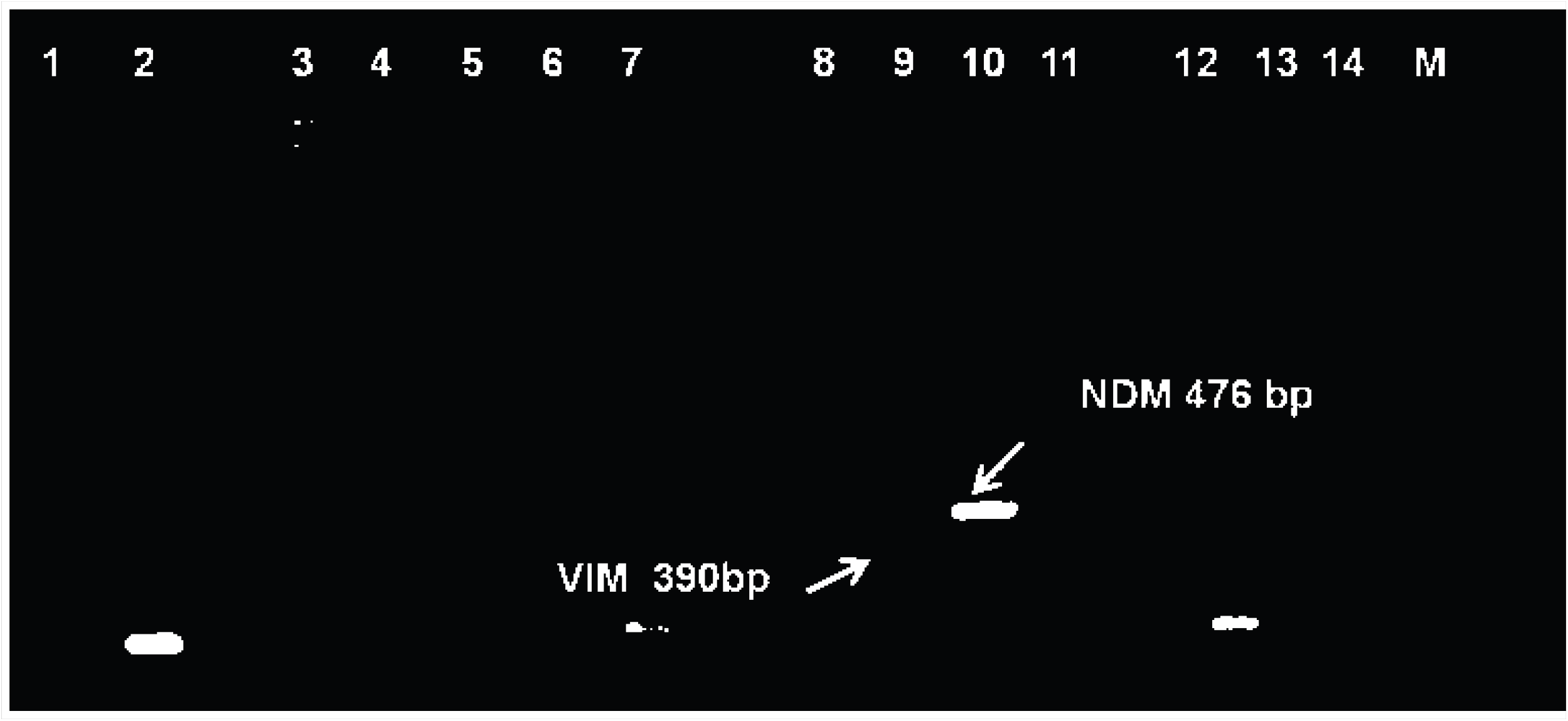
Agarose gel showing PCR amplified products of Carbapenem-resistant gene. Lane 1-ve control, Lane 2-+ control, Lane 3,4,5- NDM, Lane-6 No band, Lane 7,8 –NDM Lane 9 NDM+ VIM, Lane 10,11, 12, 13, 14-NDM. Lane M- 1kb Ladder.

**Fig 8a:**
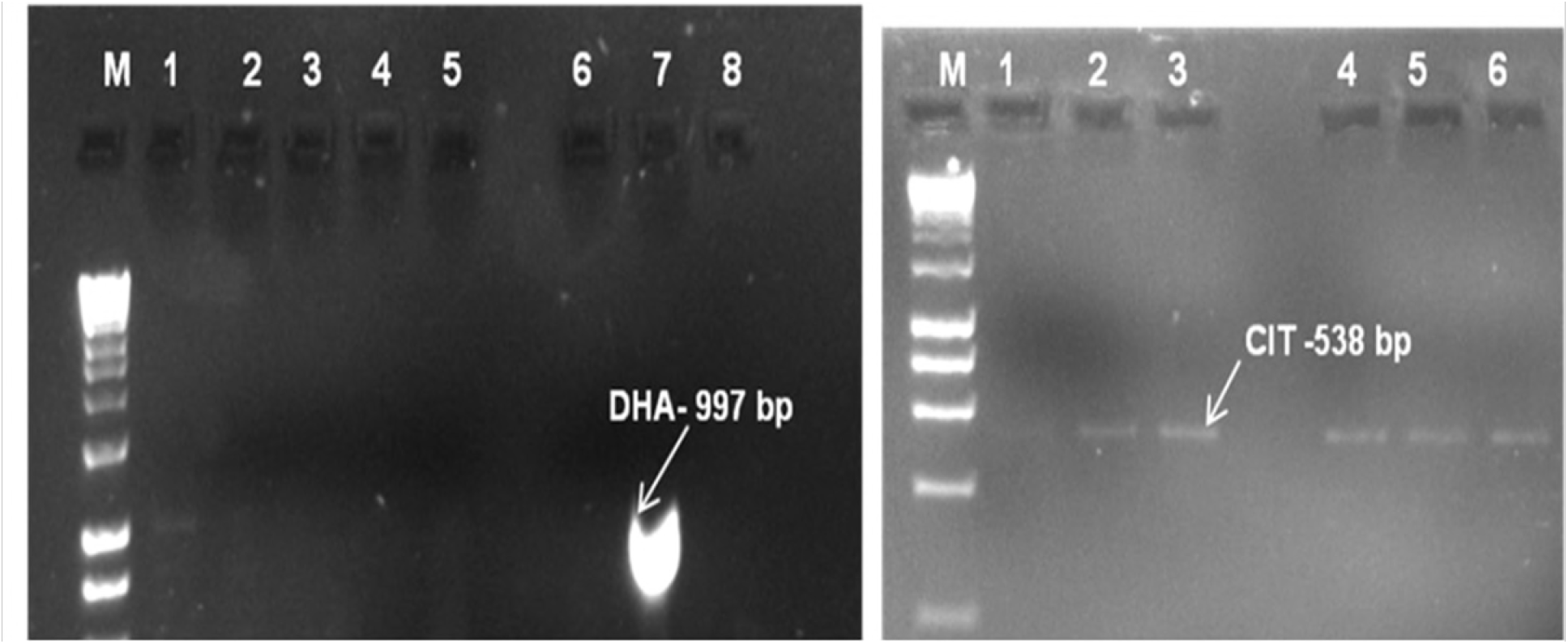
Agarose gel showing PCR amplified products of 16S rRNA methyltransferase gene. Lane M- 1kb Ladder Lane 1- - ve control, Lane 2- +ve control Lane 7- Rmt B, Lane 9- Rmt A

**Fig 8b:**
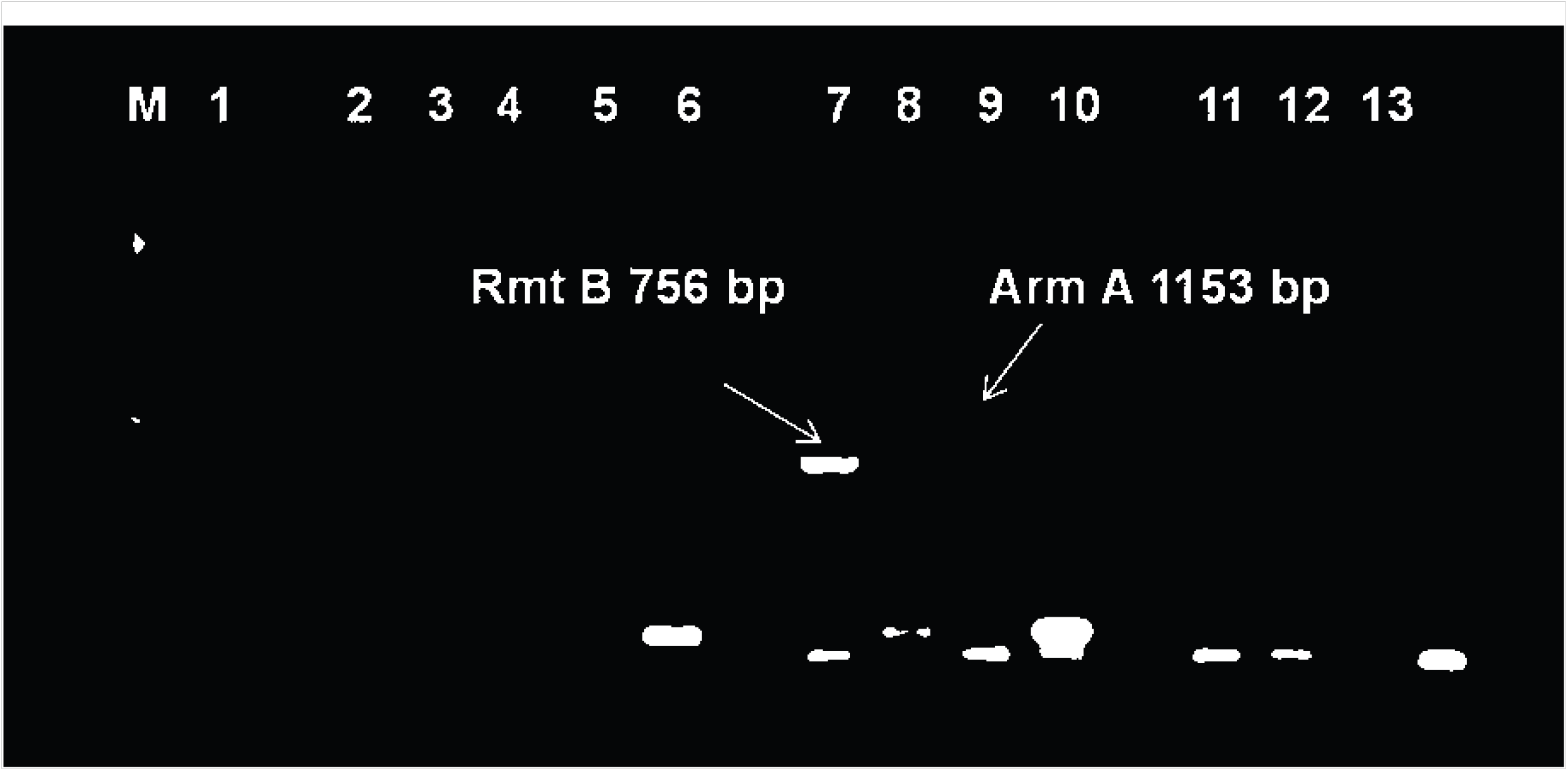
Agarose gel showing PCR amplified products of 16S rRNA methyltransferase gene. Lane M- 1kb Ladder Lane 2- Lane 4,- Rmt C, Lane 9- Rmt A, Lane 11- + control, Lane 13-ve control

**Fig 9:**
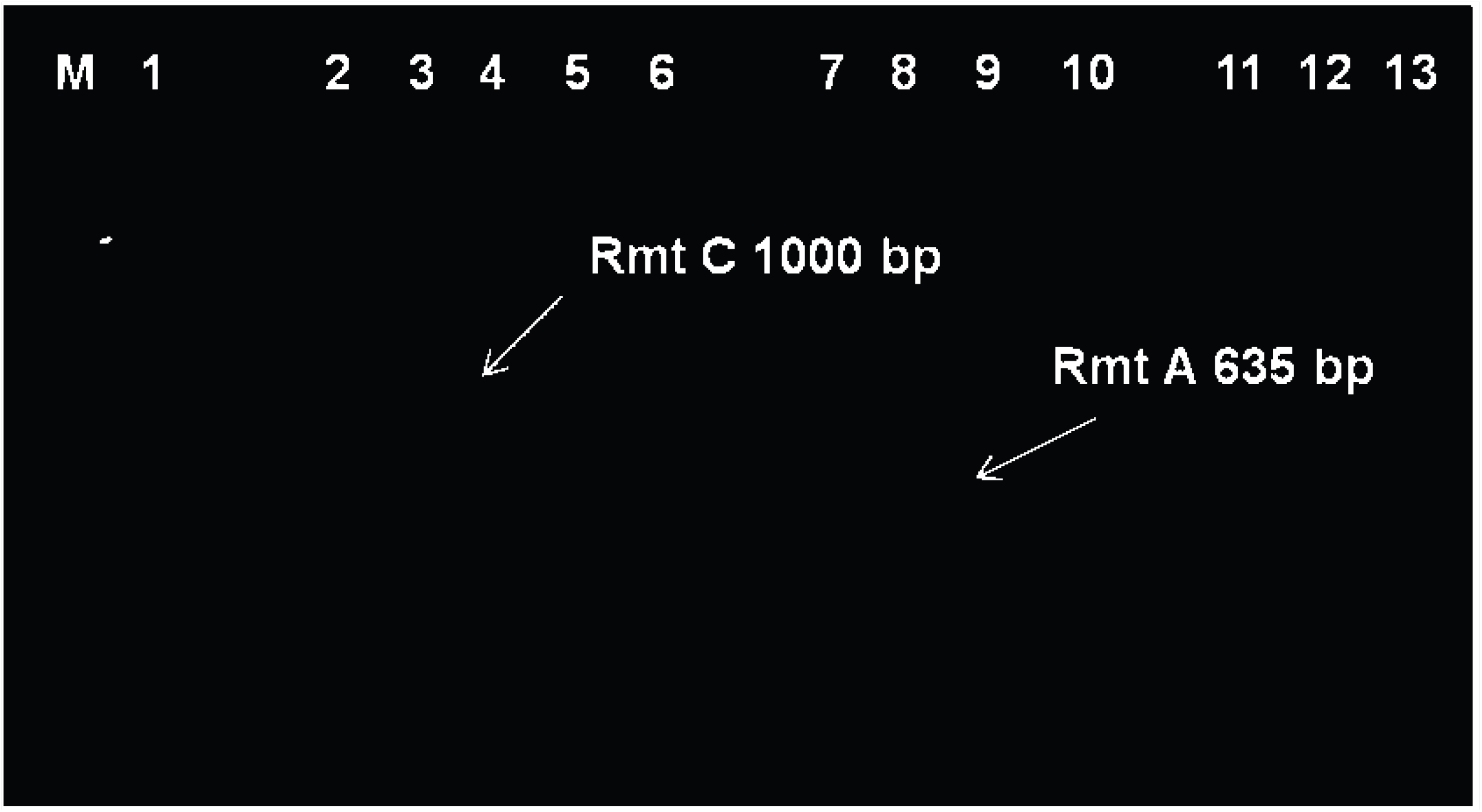
Distribution of Carbapenem resistant bacteria in uropathogen

**Fig 10:**
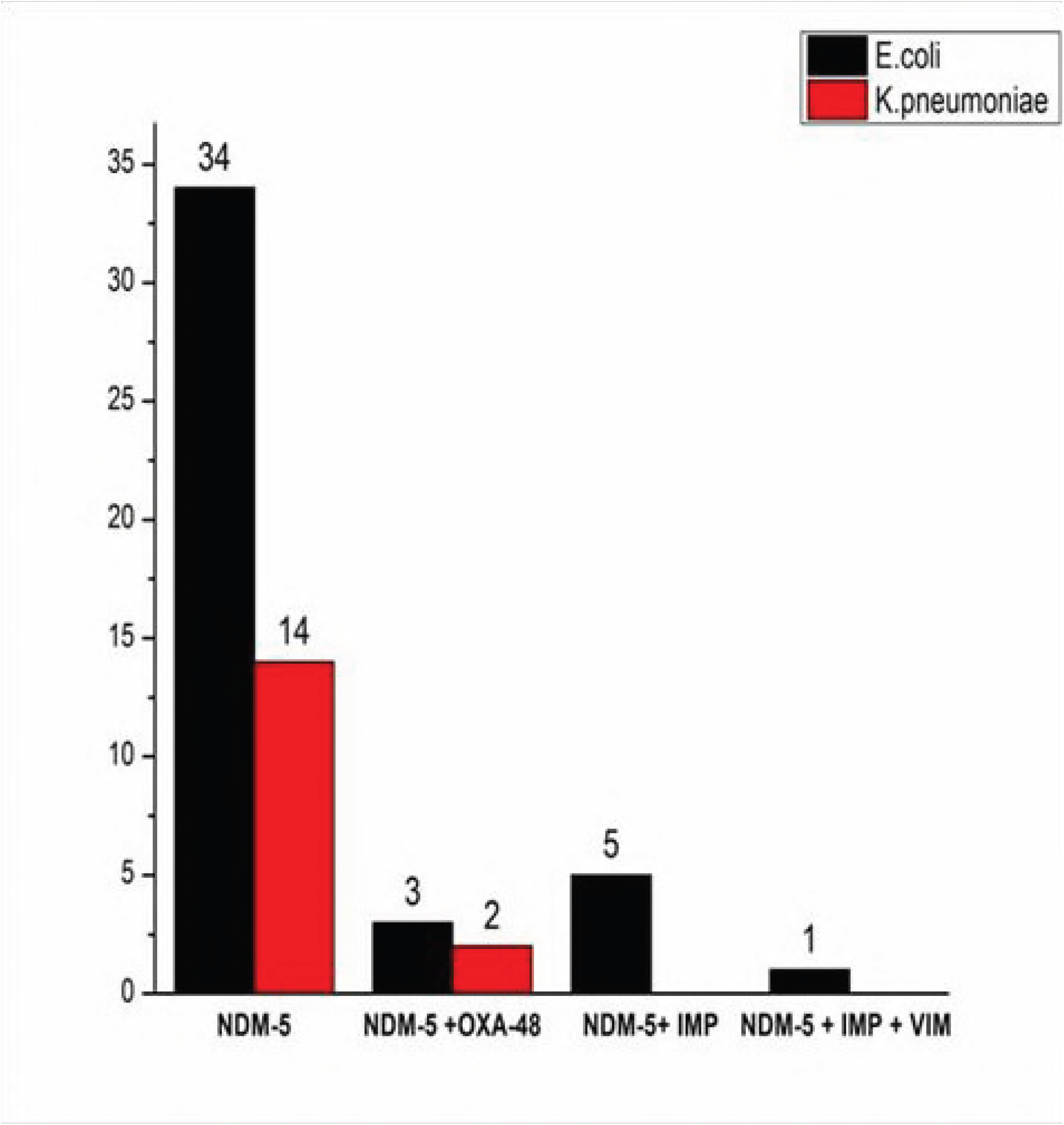
Distribution of ESBL resistant bacteria in uropathogen

**Fig 11:**
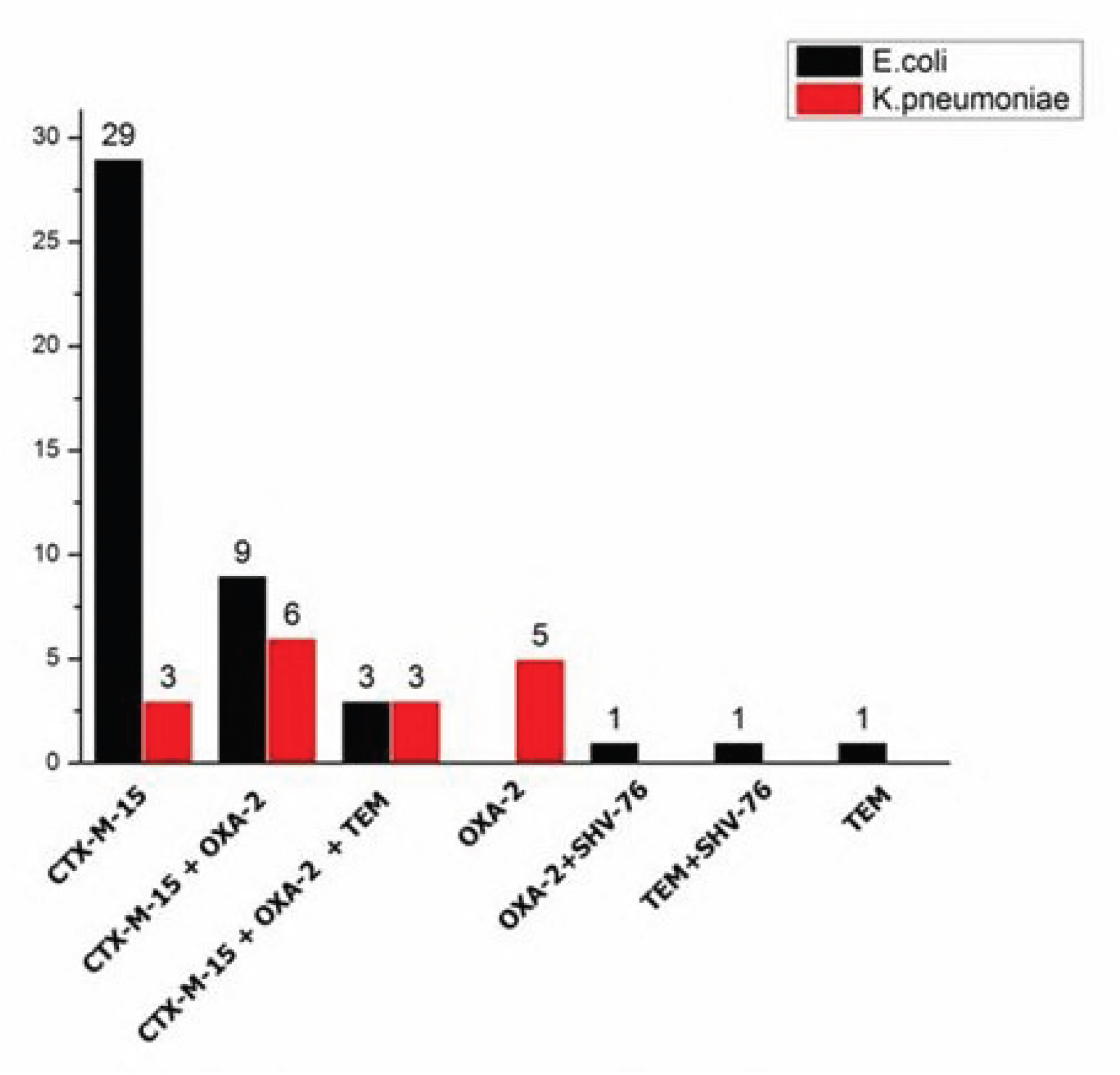
Distribution of 16S rRNA methyltransferase resistant bacteria in uropathogen

**Fig 12:**
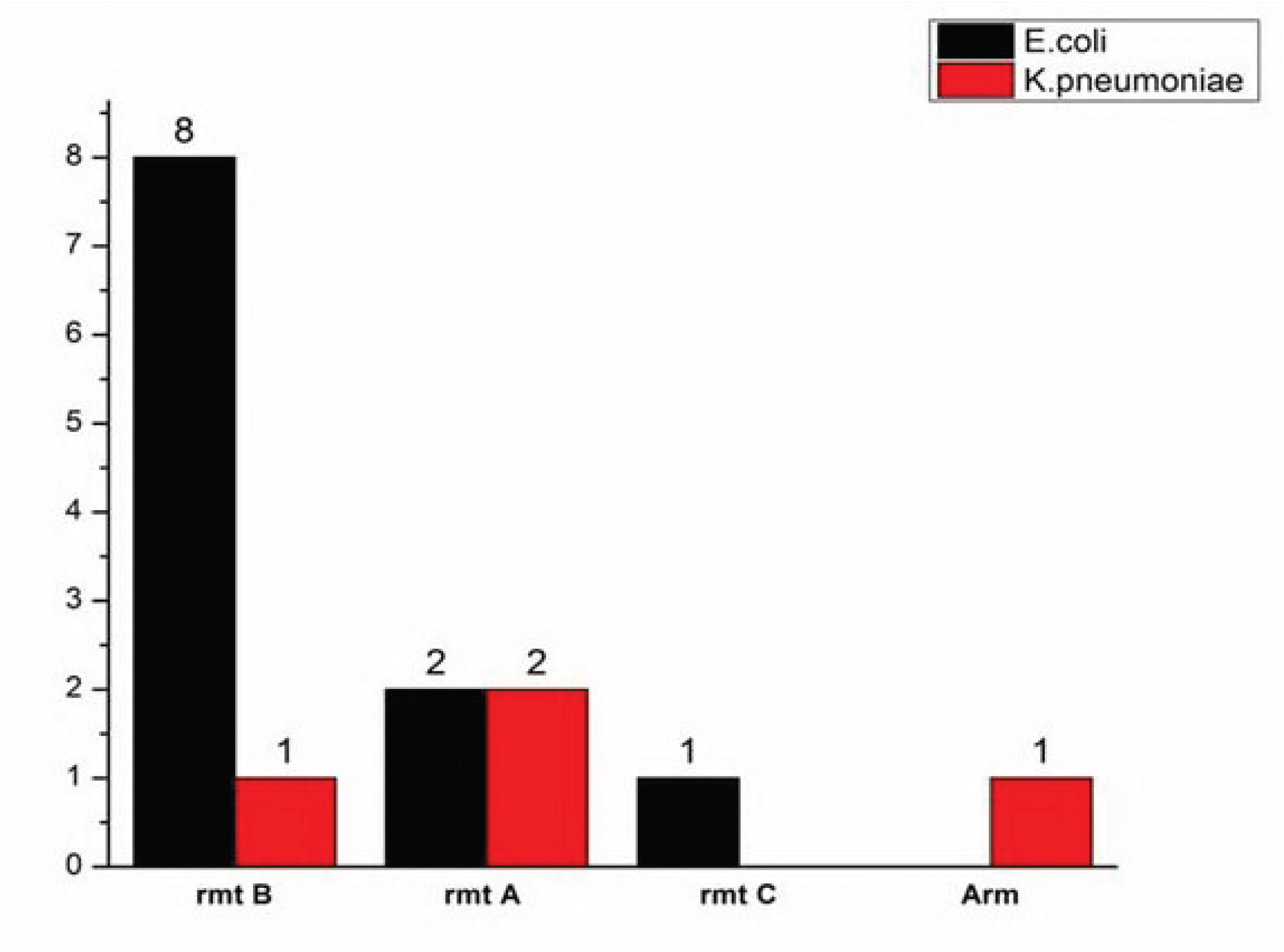
Distribution of Amp C resistant bacteria in uropathogen

CTX-M was present with another coexisting *bla* gene. From plasmid analysis, it was revealed that ESBL gene was located within the plasmid of approximately 18Kb in size. ESBL gene was found to be horizontally transferable and the resistance determinant was carried within diverse incompatibility (inc) group namely HI1, I1, FIA+FIB, FIA and Y types. Nine plasmids had an incompatibility group of FIA+FIB and Y respectively, followed by HI1 (n=5), FIA (n=4), and I1 (n=3). In the stability analysis, the above mentioned Inc type harboring ESBL genes showed progressive plasmid loss after 28 passages. This implicates the specialized adaptation of this plasmid for the survival of host under cephalosporin stress in both hospitals and in the community.

Apart from ESBL and carbapenem-resistant gene the isolates also harbored 16S rRNA methyltransferase genes (n=15) followed by AmpC genes (n=9). The most prevalent aminoglycoside resistant gene was found to be *rmt* B and the least prevalent was *rmt* C and *Arm* A (Figure 3). Amp C resistant genes included *bla*_CIT_ (n=*E.coli*-6, *Klebsiella pneumoniae*-2) and *bla*_DHA-1_ (n=*E.coli*-1) genes

### Discussion

It is reported that ESBLs in Enterobacteriaceae coexists with resistance to other antimicrobial classes and as such these organisms become multi-drug resistant hence limiting treatment options for infections. In case of infections caused by ESBL producing bacteria, carbapenems are the antibiotics of choice for the treatment [20]. However several studies have reported on the emerging resistance to carbapenem antibiotics due to the increased production of β-lactamases worldwide, which hydrolyze all β-lactam antibiotics including carbapenems. In the present study 93.65% of the ESBL producing microorgranisms harbored carbapenem-resistant genes (mainly NDM-5), most of the isolates (n=34) were obtained from community settings. Our prevalence is also much higher than data obtained in studies from Uganda where only 28.6% of carbapenemase producers were detected among ESBL producing Enterobacteriaceae.(6). Similarly, a very less prevalence of carbapenemase encoding gene was reported from Spain with a prevalence rate of 0.04% only (17). *bla*_NDM-5_ was found to be more prevalent among all carbapenemase producing genes.

The organisms harboring AmpC beta-lactamase is a major cause of therapeutic failure leaving cephalosporins inactive along with co-existing mechanism of resistance. Our study revealed a prevalence of the AmpC producing gene of 14.75% among ESBL producing Enterobacteriaceae. In our study, we found out that among all the AmpC producers, the genes producing CIT enzymes were more prevalent mainly in contrary to the study conducted by Jean et al.,2017 where most of the *E.coli* (11.7%) harbored CMY-2 producing enzyme (18).

Aminoglycosides are frequently used in combination with the ß-lactam group of antibiotics to treat severe infections in hospital patients. However, the bacterial population has developed various resistance mechanisms and very soon the therapeutic use of this drug will be limited. Acquired 16S rRNA methyltransferases which accounts for high-level and broad-spectrum aminoglycoside resistance have been reported increasingly among enterobacterial isolates in recent years, often in association with beta-lactamases, further complicating the management of infections caused by multidrug-resistant isolates (19). In our study, four different types of 16S rRNA methyltransferase genes have been characterized which are responsible for aminoglycoside resistance. Among these, *rmtB* (16.39%) was found to be the most predominant type in this part of India, on the contrary, Wangkheimayum J et al, (2017) found *Rmt C* being predominant in the Eastern part of the India (8). It is reported that 16S rRNA methyltransferases often coexist with *bla*NDM and *bla*CTX-M genes (20) (8) and our study was not an exception. The findings that ESBL producing uropathogens co-harbored carbapenem, aminoglycoside and AmpC resistant genes underscore that India as an epicenter of horizontal transfers of high-level resistance alleles between Gram-negative bacteria irrespective of community or nosocomial settings [17] [18]. Through this kind of study that has been conducted in this region, we can understand the local distribution of these ESBL resistant genes and their movement, adaptability, and propagation under antibiotic exposure in different clinical environmental conditions(23).

Since the study area shares the border of the border with countries like Nepal, Bhutan and Bangladesh a good number of patients visit these hospitals for treatment purpose. This may be one of the factors for the acquisition and spread of drug-resistant pathogens among the people of the study area.

### Conclusion

The current study revealed that the uropathogens primarily carried various types of ESBL, carbapenem, AmpC and aminoglycoside resistance genes. The emergence of multiple resistance mechanisms in these isolates makes these pathogens a major challenge in treating infections, by such pathogens as they show huge resistance to commercially available drugs. This situation is alarming and will pose a more economic burden on the people suffering from such multiple drug-resistant pathogens. In order to curb the spread of such resistant pathogens, hospitals have to come up with some antibiotic policy after regular surveillance of such type of bacterial strains.

## Declarations

### Ethical Approval

Ethical clearance for this work was obtained from the Institutional Ethical Committee, Sikkim University and consent were obtained from community participants in written form.

## Acknowledgment

The authors would like to acknowledge Rajiv Gandhi National Fellowship (UGC fellowship). The authors’ are highly grateful to the participants involved in this study.

---

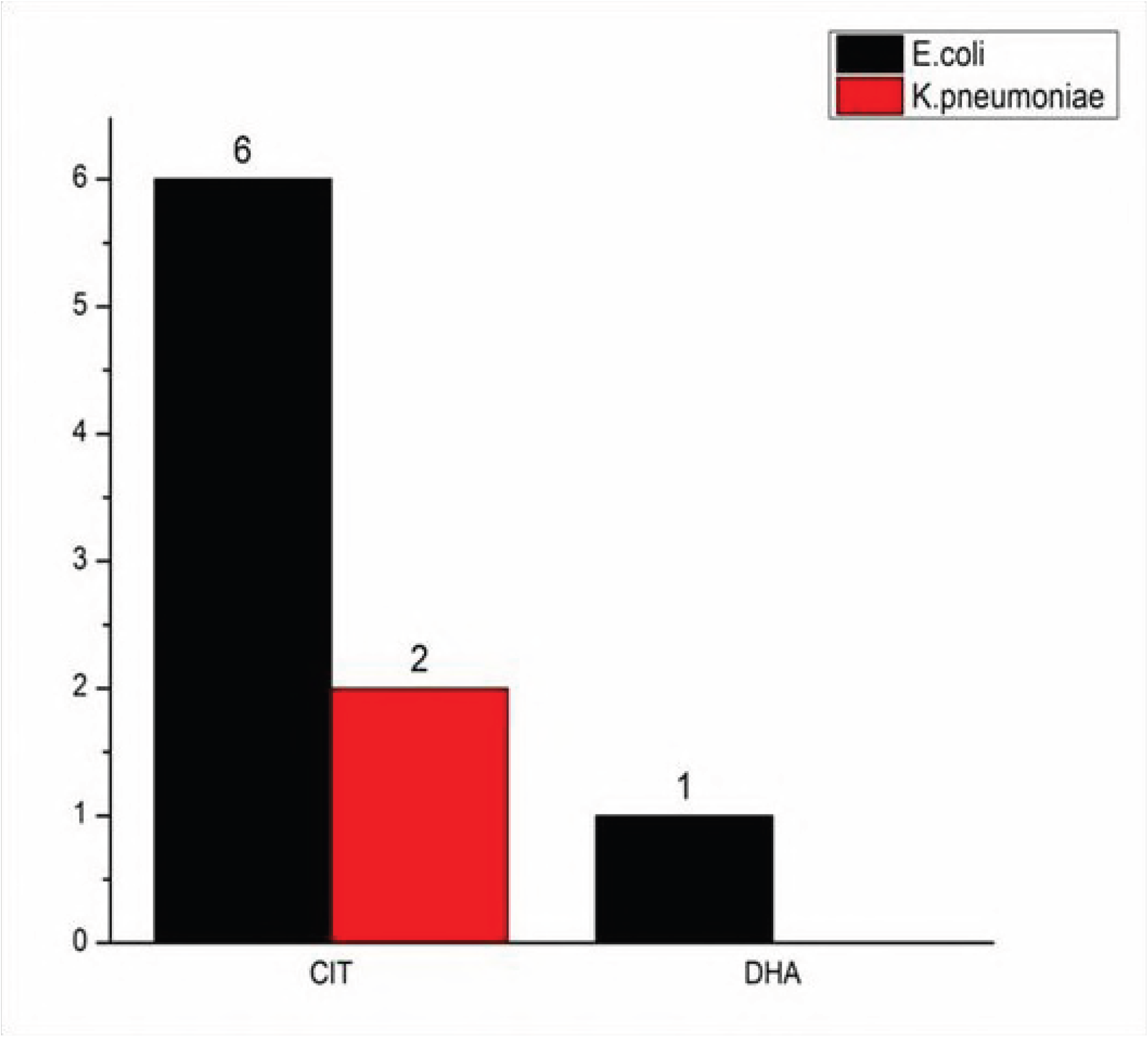

